# Understanding the Liver Under Heat Stress With Statistical Learning: A Multiomics Computational Approach

**DOI:** 10.1101/340125

**Authors:** Allen Hubbard, Xiaoke Zhang, Sara Jastrebski, Abhi Singh, Carl Schmidt

## Abstract

We present results from a pipeline developed to integrate multi-omics data in order to explore the heat stress response in the liver of the modern broiler chicken. Heat stress is a significant cause of productivity loss in the poultry industry, both in terms of increased livestock morbidity and its negative influence on average feed efficiency. This study focuses on the liver because it is an important regulator of metabolism, controlling many of the physiological processes impacted by prolonged heat stress. Using statistical learning methods, we identify genes and metabolites that may regulate the heat stress response in the liver and adaptations required to acclimate to prolonged heat stress. Our findings provide more detailed context for genomic studies and generates hypotheses about dietary interventions that can mitigate the negative influence of heat stress on the poultry industry.

**Abbreviation Term:**

RNA-seq: Ribonucleic Acid Sequencing
GWA: Genome Wide Association
SNP: Single Nucleotide Polymorphism
PCA: Principal Component Analysis
GTEX: Genotype Tissue Expression
K1: rate constant for forward reaction
K2: rate constant for reverse reaction
F6P: fructose-6-phosphate
G3P: glycerol-3-phosphate
S100Z: S100 Calcium Binding Protein Z
FBP2: Fructose-Bisphosphatase-2
NADKD1: NAD Kinase, mitochondrial
NAD: Nicotinamide Adenine Dinucleotide
NADP: Nicotinamide Adenine Dinucleotide Phoshpate
NADPH: Nicotinamide Adenine Dinucleotide Phosphate, Reduced

## Background

Obtaining biological insight from large-scale transcriptome data is challenging due to biological and technical variance. Careful experimental design can limit unwanted noise. However, when properly harnessed, heterogeneity can be used to detect biological signals that elude traditional enrichment analysis. For example, biological variation relating to a treatment response depends on many variables that are not easily controlled such as allelic or physiological variants. This fact can be informative because many compounds involved in the same process will have similar patterns of heterogeneity. This can be used to identify relationships between elements of the same pathway, even when their scales of expression and variance differ considerably, by relying on statistical learning strategies. This approach allows the combination of transcriptome and metabolome data to gain a more comprehensive biological understanding of a system. This is particularly helpful in identifying significant features from the large, complex datasets now common in multi-omics studies.

The modern broiler chicken is a fundamental source of poultry meat. It has been under strong artificial selection during the past several decades for increased breast muscle yield (Tallentire *et al.,* 2016). This is thought to be at the expense of other systems, resulting in decreased heat tolerance and increased mortality during heat stress. The relationship between the altered physiology of the broiler and susceptibility to heat stress is not fully understood, however. It is believed to involve altered appetite and preferential routing of resources to muscles tissue. Such changes are systemic, influenced by both behavior and metabolism.

One organ capable of exerting strong influence on both bird growth and thermoregulation is the liver. This organ has recently proved as a subject for studies that leverage multiomics approaches including transcriptomics and metabolomics. Such work has shed light on differentially regulated genes and metabolites. However, a systems level understanding in which fluxes in metabolites are related to gene expression, are lacking. This is partly because computational approaches exploring the totality of a biological response including gene expression and metabolite production is lacking. We combine RNA-seq (Ribonucleic Acid Sequencing) expression and metabolites from the liver to identify genes and compounds that function as biomolecules associated with heat stress. While metabolomics data identifies changes in biologically active compounds, RNA-Seq data identifies genes regulate metabolic changes. We offer a geometric interpretation for our statistical procedures, describing how they recapitulate novel biology.

Our analysis applies statistical learning approaches on metabolite and gene expression data restricts transcriptome analysis to a core module of liver enriched genes. These are determined by a definition we propose that proves more stringent than other types of relative expression analysis. Sub-setting in this fashion isolates tissue-enriched genes that reflect unique biology specific to the liver in a tissue diverse dataset. The approach of selecting tissue enriched genes and focusing on patterns of heterogeneity within such a subset provides a framework to integrate metabolite and transcriptome data. This approach of combining data from different high-throughput technologies makes it possible to identify important features of the high dimensional dataset.

Finally, extending the work of earlier GWA (genome wide association) studies that sought to model ratios of metabolites as functions of SNP’s, (single nucleotide polymorphisms) we model metabolite ratios in terms of other metabolites. The original purpose of these GWA metabolite studies was to detect genetic basis of metabolic changes (Gieger *et al.,* 2008). However, modeling ratios as function of metabolites allows detection of metabolic forks, or small network motifs where precursors are selectively routed to different metabolic fates under heat stress. The compounds used to compose triplets representing possible metabolic forks are selected from hypotheses developed through the combined k-means, random forest and PCA (principal component analysis) pipeline. A triplet is defined as a function of the form cor(A, (B/C)) where A, B and C are any combination of metabolites. Candidates for A, B and C are chosen from amino acids known to be catabolized under heat stress (Jastrebski *et al*., 2017) and sugar and fat molecules that may incorporate these molecules prioritized by our pipeline.

The combination of RNA-Seq with metabolite data identifies novel shifts in gene regulation that reflect pathway changes influencing metabolite levels. Our combined informatics strategy identifies elements under genetic control and which could be targets for selective breeding. Additionally, the identification of heat stress responsive metabolites produces candidates for feed supplementation studies.

## Methods

The heat stress response is multi-tiered and involves input from multiple tissues. At the cellular level, the heat stress response unfolds across an intricate program of organelle specific changes. Which changes are causal, and which merely correlative with underlying signal or sensing pathways, thus becomes a complex question. However, the variability associated with most basal regulators of the heat stress response should be most closely related to the variation in the heat stress response. By the transitive nature of biological communication, the introduction of noise into the signal diminishes the capacity of downstream molecules, which correlate with, but do not cause the heat stress response, to discriminate between treatment and control samples. From this perspective, the problem of identifying causal molecules from expression profile is well posed as a statistical learning problem that can be addressed through random forests. Random forests can rank candidates on their ability to correctly identify the class of samples as assigned to control or experimental treatment groups. Our approach follows sorting compounds into clusters based off of their mean expression, using k-means clustering, prior to application of the random forest algorithm and finally prioritizing these top biomolecules with PCA. After standardizing, we identify compounds most strongly associated with heat stress among liver enriched genes and metabolites. All reads were mapped to the latest NCBI release of the chicken genome and accompanying annotation, GalGal5. Mapping was done with Tophat2 and Cufflinks2, with raw counts quantification by featureCounts and differential expression accomplished with edgeR.

Biomolecules are identified and prioritized to extract pathways from whose elements triplets can be calculated. Triplets showing differential behavior selected, which demonstrate equilibrium shifts at state assumptions and thus indicate behavior of a metabolic fork.

### Geometric And Biological Consideration Of K-means Step

A goal of first leveraging k-means analysis is to build more biologically interpretable random forests, with compounds initially separated by expression level. This reflects the idea that pathways involving essential biological compounds occur across a spectrum of expression levels. Compounds with different levels of expression levels may have equally important biological roles. Separating out compounds first by this dimension prevents compounds from one expression tier crowding out those from another tier when they have similar capacities for classifying samples as control or heat stress. However, the optimal partitioning should produce clusters that are similar in explanatory power. Selecting k = 3 accomplishes this goal by distributing compounds across clusters that are as similar to one another as possible in terms of their explanatory power (Figure 2A and 2B).

### Metabolic Forks

Metabolic forks, in which ratio of metabolites represent activities of competing biological processes are an adaptation of concepts introduced by Giegier et al, in which ratio of metabolites represent biological activity of processes influence by genotype. We refer to these regulatory triplets as such, because they represent divergent fates for metabolites. Candidates for components of metabolic forks are determined via prior knowledge as compounds established in the broiler heat stress response through our previous work (Jastrebski *et al.,* 2017) and which are also biomolecules prioritized by the statistical learning components of the pipeline.

Such functions serve as a more realistic description of the biochemistry of pathway steps than simple correlations with raw data. For example, pathway steps where one enzyme regulates the forward reaction and another the reverse, the regulation through gene expression can cause relative increases in one enzyme compared to corresponding enzyme that regulates the opposing reaction. This shifts the favorability of the pathway step towards either the products or reactants, depending on the function of the enzyme. The shift in favorability towards one metabolic fate, at the expense of another, under regulation thus represents a “metabolic fork”. Having hypothesized that amino acids from catabolized proteins fuel production of sugar and fats by providing carbon backbones, we calculated “metabolic forks” that included lipids, sugar and amino acids. P-values are determined from the interaction term of the resulting linear model of the metabolic fork, signifying a difference in slope between control and experimental conditions.

## Results

## Discussion

Our complete pipeline, which combines statistical learning techniques with hypothesis-free modeling of metabolite ratios, is able to propose novel hypotheses while recapitulating significant known biology from the liver metabolome and transcriptome (Figure 1). Importantly, this perspective identifies changes in compounds with roles across organelles that are increasingly thought to have important functions in the heat stress response.

**Figure 1:**
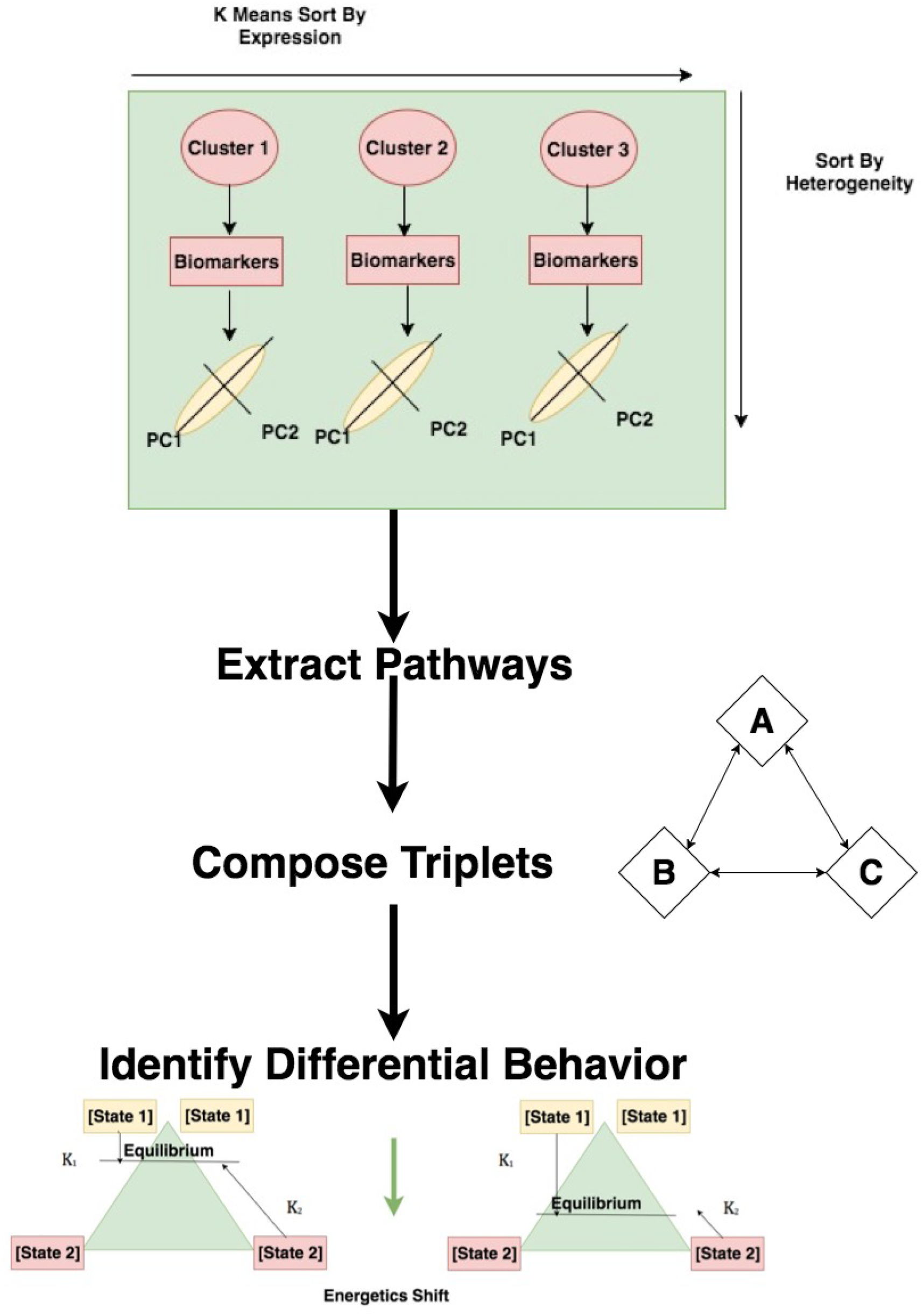
Total pipeline, from data analysis to identifying hypothetical mechanisms.

Much interesting biology, for example, relates to changes in the cell membrane. There are widespread shifts in levels of constituent lipids, for example.

The exact mechanisms by which these shifts occur remain unclear, but accumulating evidence suggests these changes in the cell membrane exert important downstream effects on heat stress responsive genes and metabolites. At least some of these may be driven by dietary changes. One such example is the essential fat linoleic acid, which is a precursor to arachidonic acid and emerges as a strong heat stress associated biomolecule and whose detected levels are lower under heat stress. The compound also correlates with two principal components among the heat stress associated biomolecules among its cluster (Additional File 1, Tables 2 and 3). Downstream arachidonic acid derivatives are similarly decreased, many of which have roles in inflammatory response.

Other biomolecules prioritized through correlation with the same principal component include other lipids, related to signaling and fatty acid oxidation – such as adipoylcarnitine and the taurine related endocannabinoids N-oleoyl taurine and N-Stearoyl taurine (Additional File 1, Tables 2 and 3). These compounds represent a possible intersection between signaling lipids and sulfur metabolism via coupling with taurine. All of these compounds occur at lower concentrations under heat stress. While the mechanisms of such regulation remain unclear, there is much evidence that suggests lipid changes influence cell state and, potentially, bird metabolism. Lipid changes, in fact, are increasingly recognized as potential regulators of heat stress at a fundamental level (Balogh *et al.,* 2013).

Recent studies have focused on nuances of the heat stress response by revising the model that it is primarily triggered by the presence of unfolded proteins (Hoffman, 2007). For example, lipids in the cell membrane may detect membrane disorder and other physical consequences of heat stress and trigger signal cascades (Balogh *et al.,* 2013). The evolutionary value of using a thermo-sensitive organelle such as the cell membrane to refine the heat stress response is the advantage of being able to regulate homeostasis through sensitive adjustments that have meaningful influences on cell fate (Balogh *et al.,* 2013). The inflammatory response may be a significant component in the transition from heat stress to heat stroke.

## Heat Stress, Membranes and Lipids

The sophisticated signaling environment created by the cell membrane is comprised of a diverse set of lipids and proteins. Among these is an abundance of sphingolipids that form rafts in the membrane and possess important signaling roles (Simons and Ikonen, 1997). The organization of the cell membrane is intricate and becomes dynamic under stress response. Important structural changes occur through interactions with membrane proteins, the gating of which possess thermal sensitivity (Torok *et al.,* 2014). Additionally, heat causes changes in physical attributes such as diffusion and dimerization rates. Measurements suggest these characteristics change in a predictable fashion during even mild heat stress events (Torok *et al.,* 2014). Thus, the cell membrane is well equipped to sense relative temperature changes.

Not surprisingly, among the compounds prioritized by our pipeline are many lipids. These shifts suggest mixture of changes in compounds with signaling and structural roles. Alterations in lipid content are important in thermal shifts associated with both heat stress and extreme cold. For example, a key adaptation to cold is the increase in membrane fluidity mediated by elevating the fraction of cis-unsaturated fatty-acyl groups in membrane lipids (Vigh *et al.,* 1998). Alternatively, during episodes of heat stress mechanisms to endure temperature shifts focus generally on maintaining the integrity of the cellular processes and such pathways can be causally regulated by changes in cell membrane disorder (Vigh *et al.,* 1998). Regulation of heat shock factors can be influenced by addition of saturated and unsaturated fatty acids, with the former inducing expression and the latter suppressing it (Carratu *et al.,* 1996).

The possibility that the qualities of the cellular membrane make it an ideal substrate in which to store ‘memory’ or serve as a ‘control center’ for a physiological response in terms of the composition of density and sensors could be extremely interesting biologically. This could prove extremely important in terms of identifying mechanistic regulators of the general response. Indeed, changes in membrane fluidity induced via alcohols triggers systemic responses paralleling those caused by heat stress, albeit in the absence of any thermal activation. Such changes include hyperpolarization of the mitochondrial membrane (Balogh *et al.,* 2005). Such experimental work confirms the role of lipids from a regulatory perspective and the influence of the heat stress response across organelles.

Among the cell membrane lipids influenced by heat stress and which are prioritized among their respective clusters is a number of sphingomyelin species. These are substantially down regulated under heat stress, and emerge as strong classifiers in clusters one and two. This is a potentially significant observation in the context that sphingolipids are up-regulated in the early phases of acute heat stress in studies of yeast (Jenkins *et al.,* 1997). Many of these sphingomyelin species correlate with principal components among their clusters that include the downregulated inflammatory arachidonic acid derivatives (Additional File 1, Tables 7 and 8). Their general attenuation may be an important aspect of physiological adaptation to the long term heat stress experienced by the birds, with the pattern of hetereogeneity in their levels indicative of bird acclimatization.

## Anti-Oxidants and Energy Burden

Heat stress entails a number of challenges that endanger cell function and which must be addressed in order to preserve homeostasis. The management and deployment of protective systems can be quite independent from the initial sensory capacity of the cell membrane. These, for example, can respond to states of cellular stress that could be ongoing in a state of heat stress. Such pathways are essential to the heat stress response, as they relate to management of general consequences of oxidative damage. Several precursors of anti-oxidants, as well as such compounds themselves, are identified as strong classifiers of treatment assignment within each cluster. These compounds manage the effects of toxic intermediates resulting from increased energy production, mitigating their ability to damage DNA or organelles. Their production may exploit the carbon backbones of amino acids released by catabolized protein.

Not surprisingly, given the relationship between oxidation and energy production, some of the classifiers suggest changes in mitochondrial activity. Even slight changes in cell resting state can have dramatic changes on the production of reactive oxygen species and the behavior of the mitochondria (Akbarian, 2016). Molecules associated with mitochondrial performance are computationally recognized as potential biomarkers of the heat stress response. This suggests that mitochondrial conditions are closely related to general heat stress, and that the cell adjusts antioxidant levels accordingly.

At the same time that sugars and other energy-related metabolites show upregulation, an important class of lipids involved in the carnitine shuttle system that transports fatty acids to the mitochondria shows consistent downregulation. These carnitine species (linoleoylcarnitine, stearoylcarnitine, adipoylcarnitine) are identified as strong heat stress associated biomolecules among their clusters and correlate strongly with resulting principal components (Additional File 1, Tables 1, 2 and 3). Such patterns suggest sweeping downregulation of fatty acid oxidation pathways, as metabolism is increasingly driven by gluconeogenesis. Transcriptome changes support a coordinated shift in lipid and sugar management (Jastrebski *et al.,* 2017).

Genes that correlate most highly with the principal components that emerge from the k-means cluster containing gluconeogenesis biomolecules include Neurotensin Receptor 1 (NTSR1) (Additional File 1, Tables 4 and 5). However, the correlation of this transcript with the first principal component, at .48, is relatively weak compared to the main metabolites associated with gluconeogenesis, i.e. .48. Glucosamine-6-phosphate and glucose-6-phosphate have correlations of .89 and .91, for comparison. However, NTSR1 correlates much more strongly with the second principal component, at .75. NTSR1 is a neurotensin receptor, and is associated with appetite regulation and blood sugar levels. Another strong classifying gene associated with this cluster is the methyltransferase METTL7A, though it only correlates significantly with the second principal component (.703) (Additional File 1, Table 5). Upregulated NTSR1 may be important to maintaining blood sugar homeostasis in the presence of enhanced gluconeogenesis, whereas METTL7A may be manage sulfur metabolism associated with enhanced antioxidant production. In addition to

This shift towards gluconeogenesis is supported strongly from a mechanistic standpoint by the metabolic fork (Figure 10). The metabolic fork provides evidence of large-scale redirection of carbon resources released from the catabolized glycine. This complements the statistical learning pipeline, which prioritizes biomolecules without determining whether they are causal or merely collinear to biological changes. The detection of a potential metabolic fork also provides kinetic information about biochemical pathways that may be consequences of gene expression changes. Directionality can be inferred through prior knowledge. For example, the metabolic fork suggests that carbon released from catabolized glycine is preferentially routed to F6P through gluconeogenesis, as opposed to G3P, under heat stress. This is supported by increased transcription of the rate-limiting gene controlling this step, FBP2, under heat stress. The ability to identify concrete mechanisms which exhibit differential behavior under heat stress, and which are consistent with transcriptome changes, makes it possible to complement purely correlation-based strategies with mechanistic hypotheses.

## Metabolic Forks Resulting From Gene Regulation

One of the top differentially regulated triplets contains two compounds prioritized through PCA on top biomolecules on a k-means cluster. This is consistent with gene important expression changes, such as those involving FBP2. The three members of the triplet span gluconeogenesis (fructose-6-phosphate), glyceroneogenesis (glycerol-3-phosphate) and amino acid catabolism (glycine). Pairwise correlations between each node are provided on the edge corresponding edge. A proposed mechanism for the observed pattern is that catabolized glycine is preferentially shunted towards gluconeogenesis under heat stress, thus contributing to F6P production. Increasingly fueled by carbon backbones provided by amino acids from catabolized proteins, gluconeogenesis decouples from glyceroneogenesis under heat stress.

The ratio of G3P to glycine represents the tendency of catabolized amino acids to become backbones for fats, as opposed to sugars. This changes as a function of increased demand for sugar under heat stress and is corroborated by increase in the gene Fructose-Bisphosphatase-2 (FBP2) encoding the rate-limiting gene for gluconeogenesis.

## Summary and Future Work

Interest in the heat stress response is broad, stretching from plant physiology to human clinical research, with insights potentially applicable across taxa due to the deep conservation of cell signaling pathways. Next generation sequencing technologies provide new experimental perspectives from which to explore such systems. During the past several years, the advent of next generation sequencing tools has produced a deluge of data. However, methods to process that data have been lacking. Combining the information from multiomics and multi organ datasets compounds this challenge. The capacity to link patterns of heterogeneity to pathway importance is an approach that can ease the burden of prioritizing compounds in such a setting. Here, we do so and leverage a combination of relative tissue enrichment and statistical learning approaches to prioritize compounds based on their ability to identify samples as belonging to heat stress or control conditions. We demonstrate signatures of the heat stress response across several important systems. Importantly, this is a very general strategy that works with any type of continuous data, rendering it applicable to both metabolome and transcriptome data and flexible enough to accommodate future “-omics” data. From both types of “-omics” data, we identify a diverse range of important mechanisms that may be influencing shifting biology.

We have leveraged a flexible method of analysis to process a complex dataset and identify important components of the heat stress response. While recapitulating known biology, our analysis also proposes new hypotheses about heat stress regulation that relates to systems controlled by a diverse range of organelles. These can be explored through future experimentation. Additionally, the metabolic fingerprint of heat stress provides candidates for feed supplementation studies. Thus, this study proposes a general workflow to integrate high dimensional, complex datasets in order to yield testable hypotheses about biology.

**Figure 2A and 2B:**
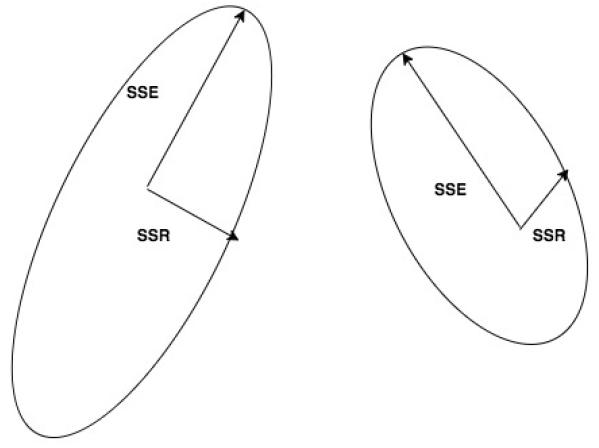
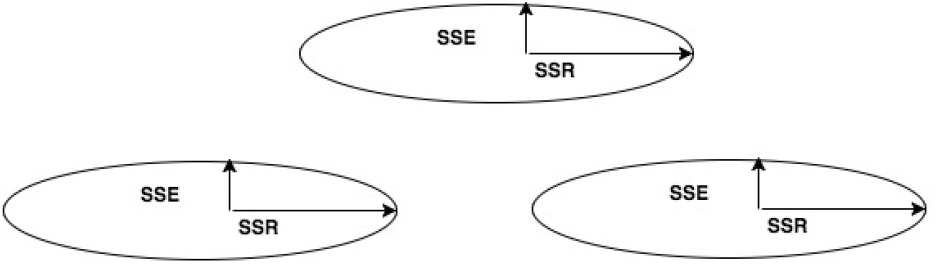
Example of possible models around specific cluster with different k-means selection, illustrating more uniform clustering results with k = 3 (2B) compared to k = 2 (2A).

**Figure 3:**
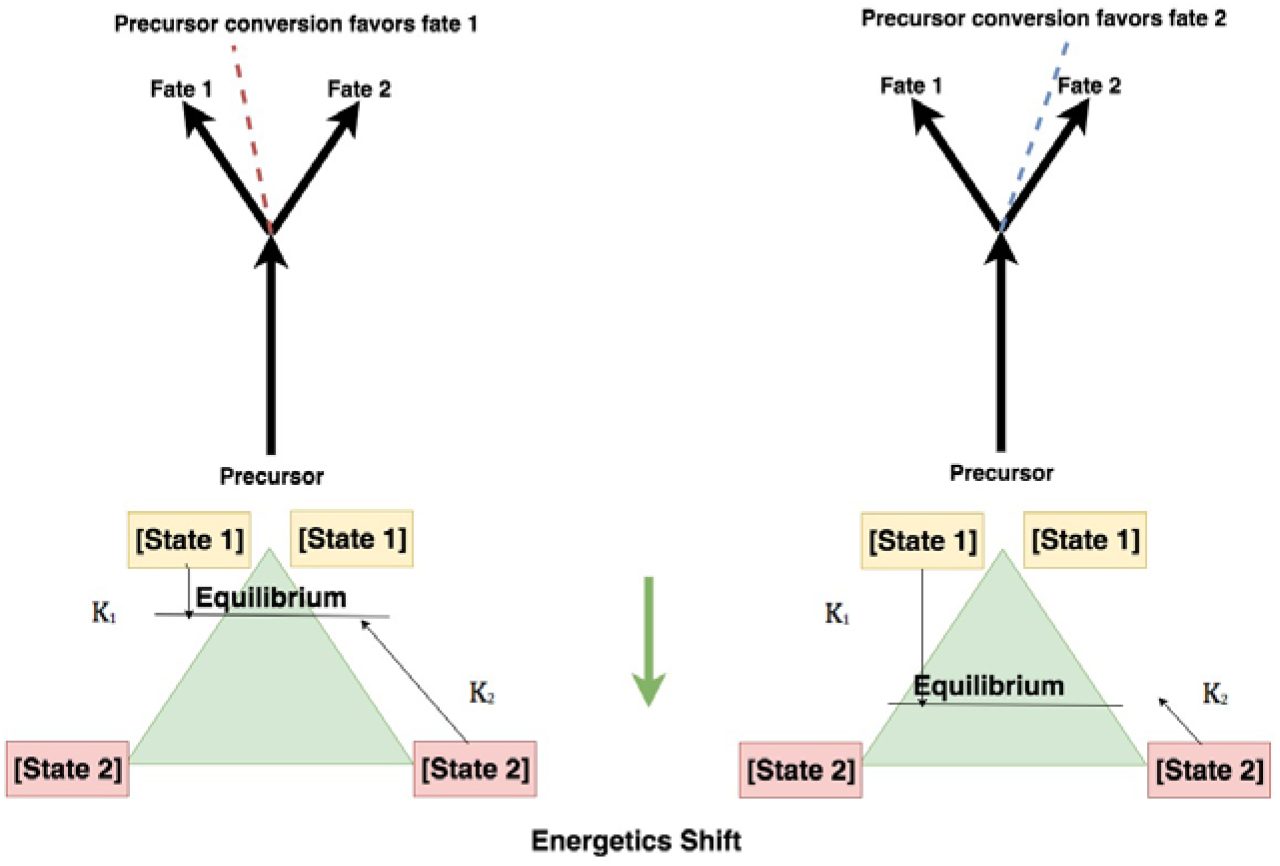
Under changes in gene expression that alter levels of the regulating enzymes, precursors are preferentially routed to one metabolic fate over another. Shifts in the ratio between metabolites representing fate 1 or fate 2 may represent shifts in biology.

**Figure 4:**
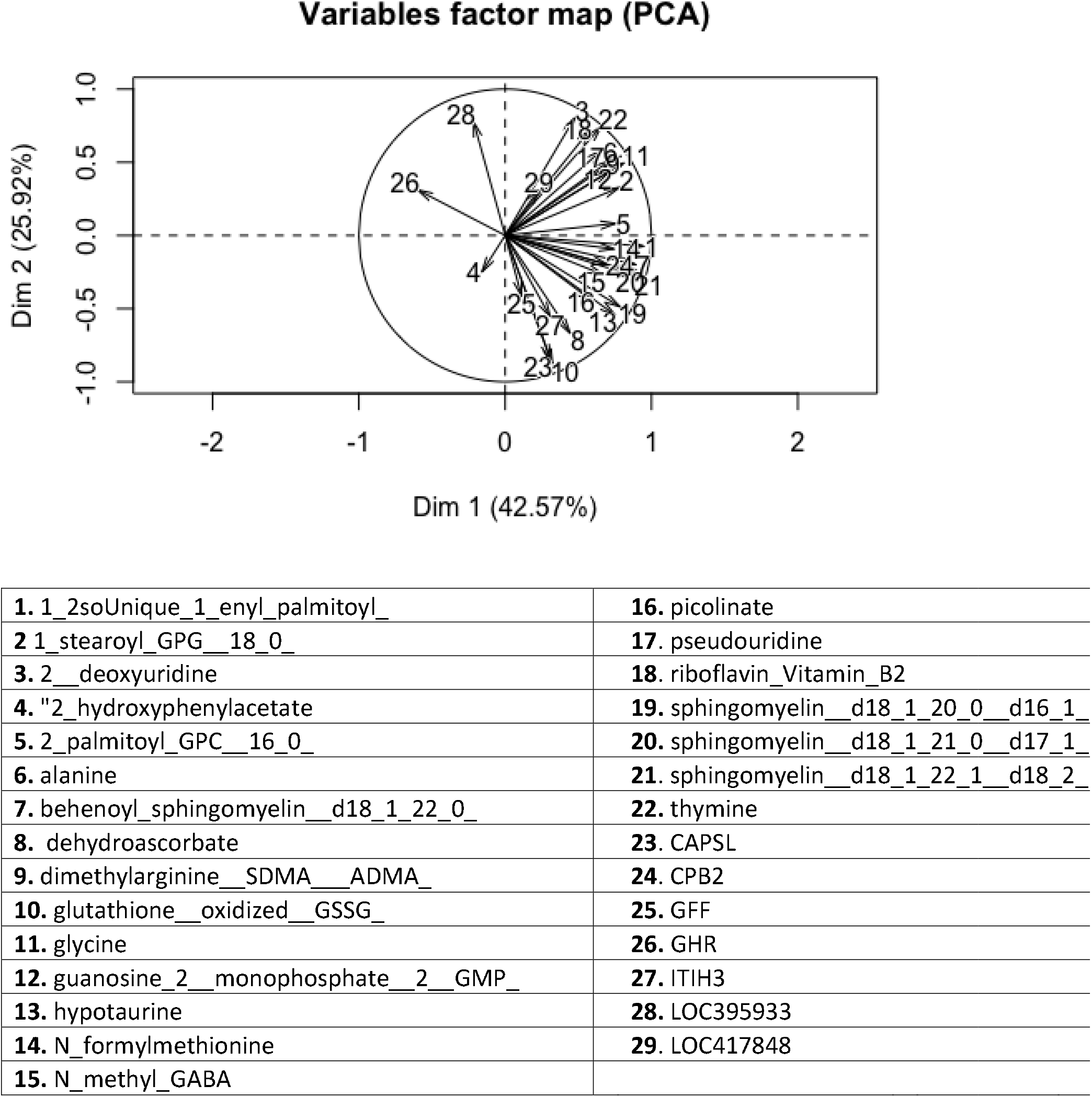
Biplot of PCA analysis on correlation matrix of the top biomolecules in cluster 1. See keys to match numbers to compounds.

**Figure 5:**
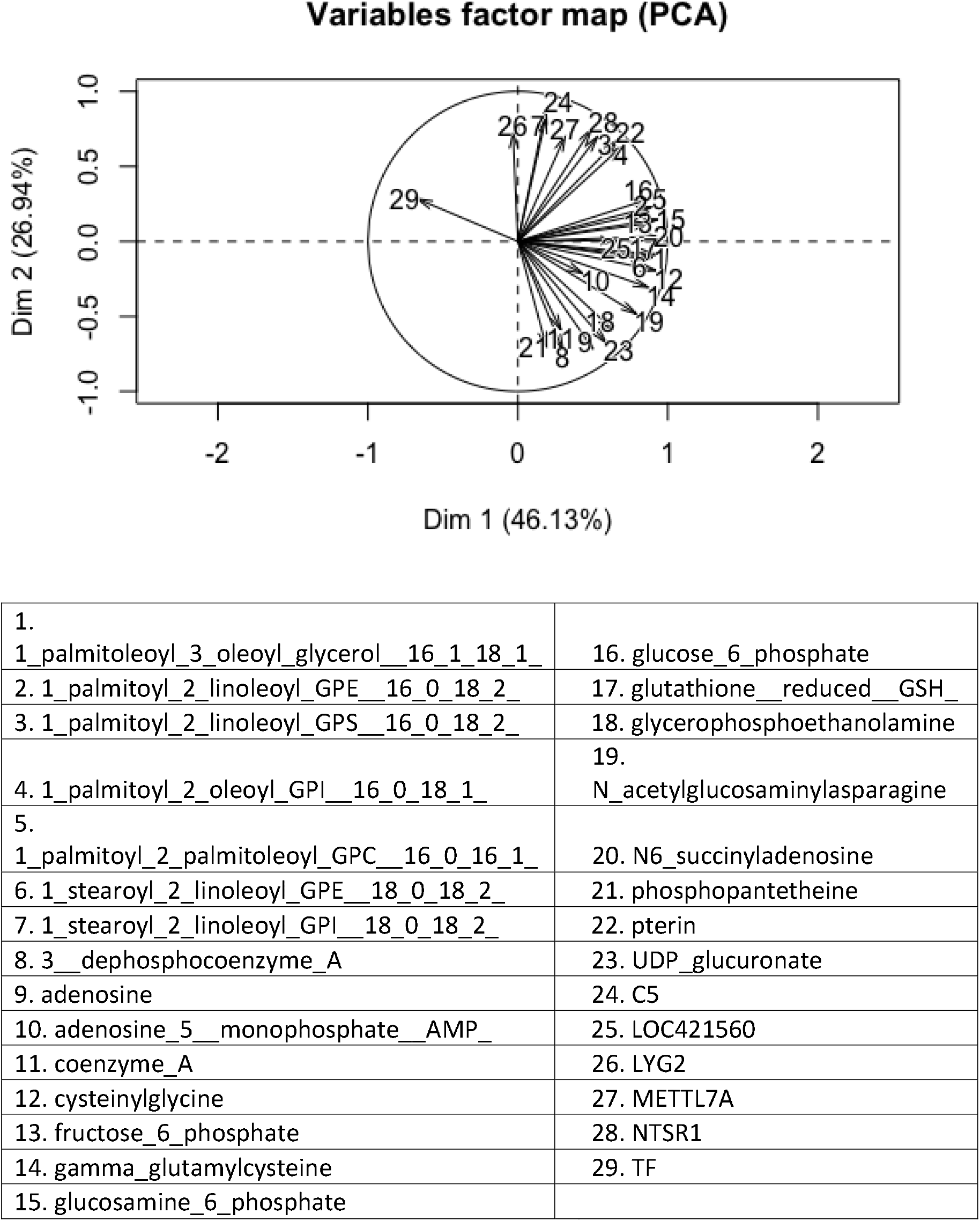
Biplot of PCA analysis on correlation matrix of the top biomolecules in cluster 2. See keys to match numbers to compounds.

**Figure 6:**
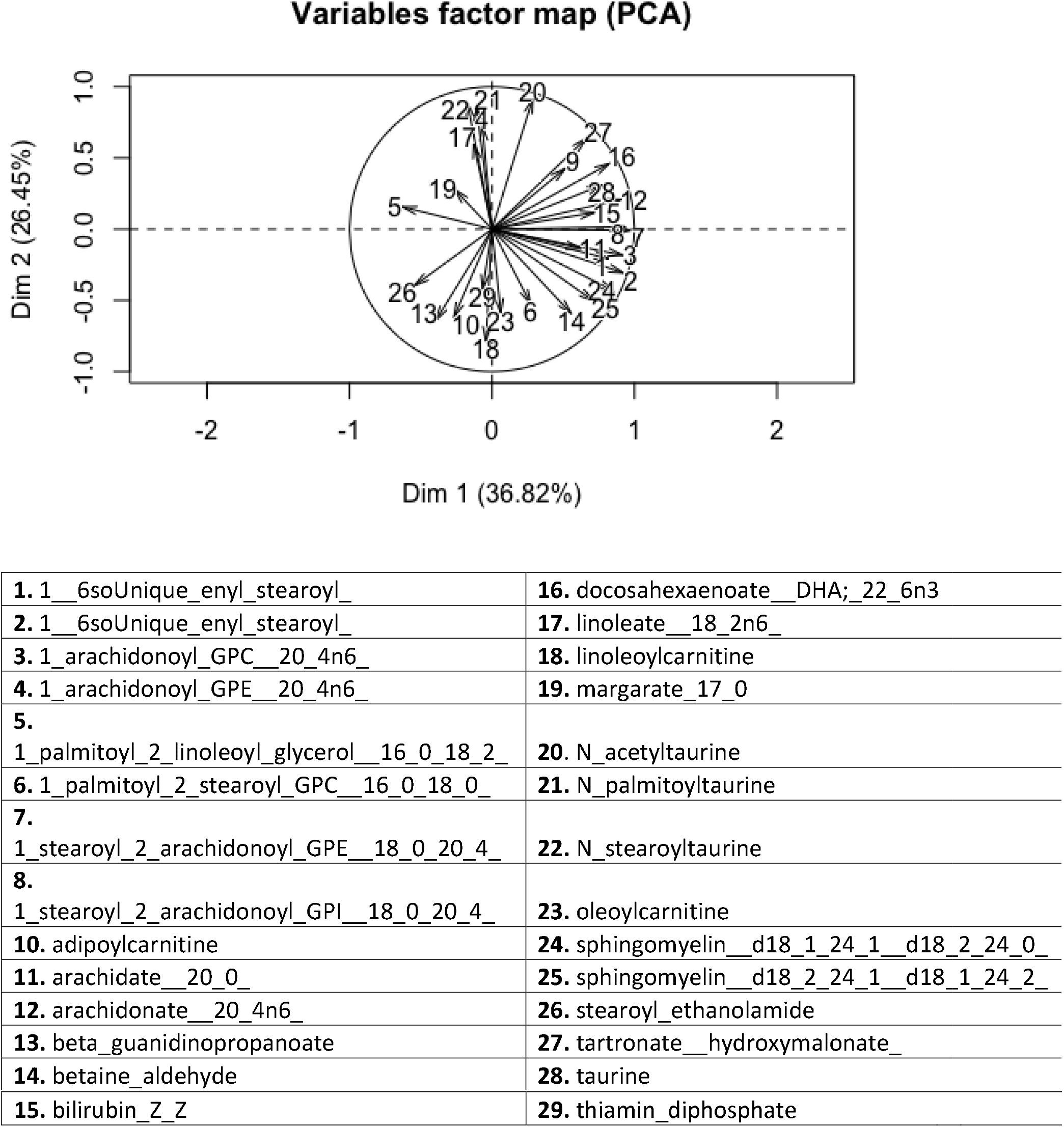
Biplot of PCA analysis on correlation matrix of the top biomolecules in cluster 3. See keys to match numbers to compounds.

**Figure 7.**
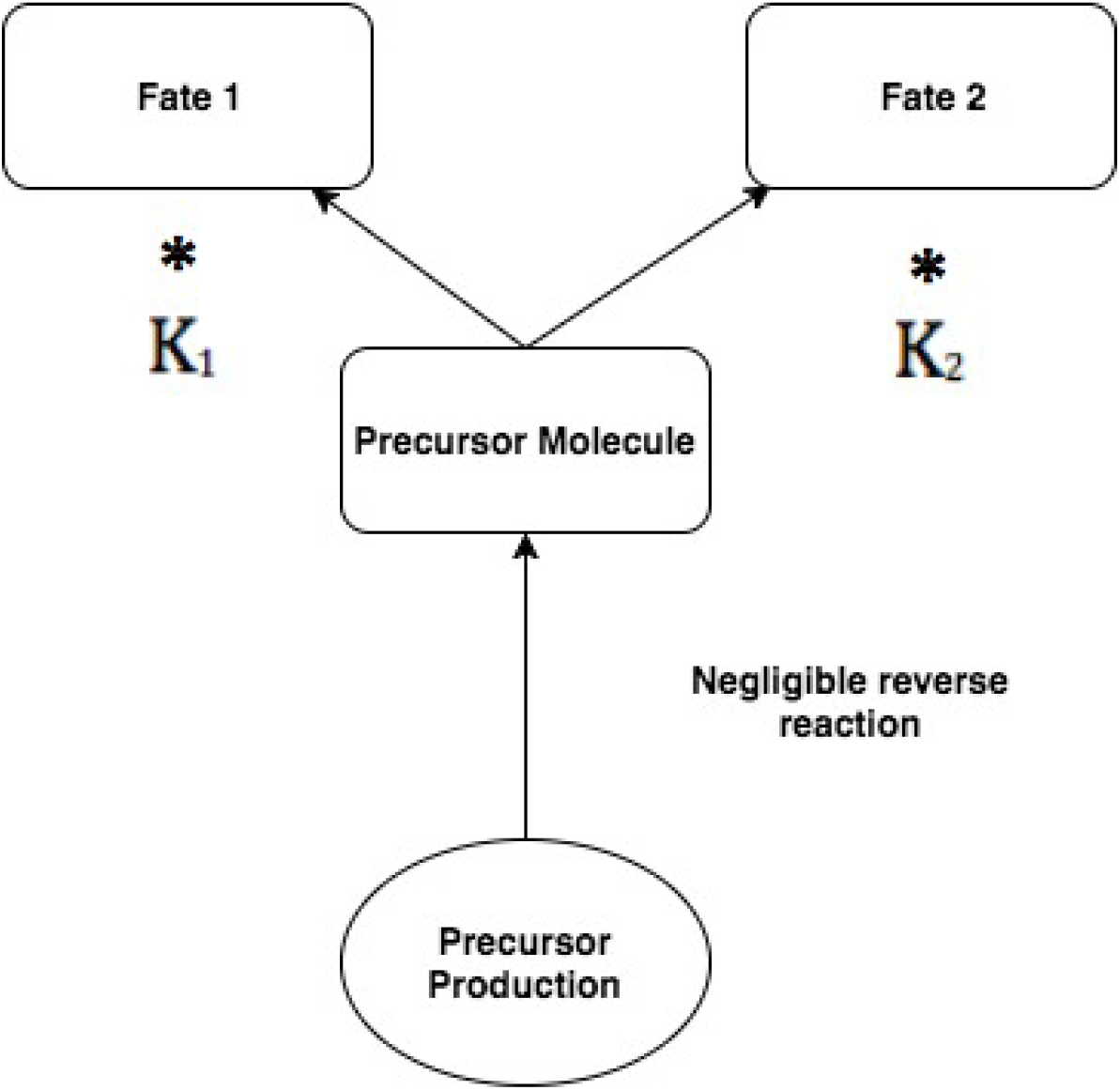
Illustration of the components of a metabolic fork.

**Figure 8:**
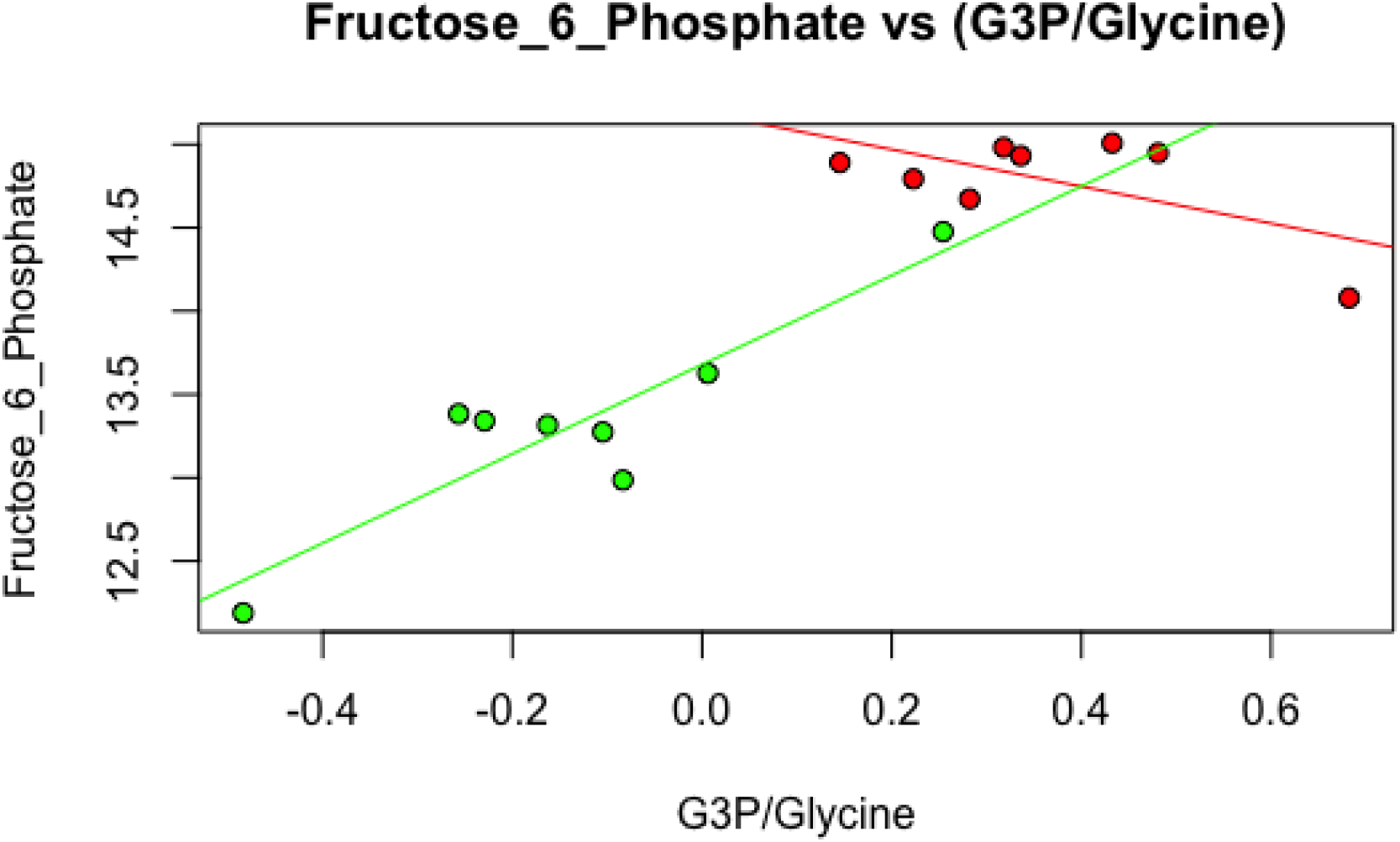
Fructose-6-phosphate (F6P), a sugar produced by gluconeogenesis, as a function of the ratio of glycerol-3-phosphate (G3P) and glycine. All values are log transformed. P-value of interaction term = 0.0005.

